# Dynamic regulation of anterior-posterior patterning genes in living *Drosophila* embryos

**DOI:** 10.1101/2020.11.25.395277

**Authors:** Takashi Fukaya

**Author notes:** Correspondence should be addressed to T.F.

## Abstract

Expression of the gap and pair-rule genes plays an essential role in body segmentation during *Drosophila* embryogenesis [1–5]. However, it remains unclear how precise expression patterns of these key developmental genes arise from stochastic transcriptional activation at the single cell level. Here, I employed genome editing and live imaging approaches to comprehensively visualize regulation of the gap and pair-rule genes at the endogenous loci. Quantitative image analysis revealed that the total duration of active transcription (transcription period) is a major determinant of spatial patterning of gene expression in early embryos. The length of transcription period is regulated by the continuity of bursting activities in individual nuclei, with core expression domain producing more bursts than boundary region. Each gene exhibits distinct rate of nascent RNA production during transcriptional bursting, which contributes to gene-to-gene variability in the total output. I also provide evidence for “enhancer competition”, wherein a distal weak enhancer interferes with transcriptional activation by a strong proximal enhancer to downregulate the length of transcription period without changing the transcription rate. Analysis of endogenous *hunchback* (*hb*) locus revealed that the removal of distal shadow enhancer induces strong ectopic transcriptional activation, which suppresses refinement of broad expression domain into narrower stripe pattern at the anterior part of embryos. This study provides key insights into the link between transcriptional bursting, enhancer-promoter interaction and spatiotemporal patterning of gene expression during animal development.

## Results and discussion

Zygotic transcription of the gap and pair-rule genes plays an essential role in body segmentation of *Drosophila* embryos [1–5]. Each gene produces distinct spatial pattern with sharp ON/OFF boundary from crude gradients of maternal morphogens [6]. Positional information encoded by anterior-posterior (AP) pattering genes is not static, rather it undergoes dynamic spatial shifting over time [7], which is thought to facilitate time-dependent changes in the distribution of regulatory gradients within an embryo. Importantly, recent advent of quantitative imaging methods revealed that transcription is a stochastic process that consists of successive bursts of *de novo* RNA synthesis in various species including *Drosophila* [8–16]. Nonetheless, spatiotemporal patterning of developmental genes is highly reproducible within a population [17]. Thus, question remains to be addressed how precise expression patterns arise from stochastic transcription in developing embryos.

To directly visualize transcription dynamics of AP pattering genes at the endogenous loci, I employed genome editing and quantitative live imaging approaches [18–20]. The sequence cassette containing 24x MS2 and SV40 poly(A) signal was inserted into the 3′ untranslated region (UTR) of target genes using CRISPR/Cas9 (Figure 1A). First, MS2-tagged strains were obtained for four major gap genes, *giant* (*gt*), *hunchback* (*hb*), *Krüppel* (*Kr*) and *knirps* (*kni*) (Figure 1B and Figure S1A). The length of inserted MS2 cassette is same for all genes, permitting direct comparison of transcription activities of different genes irrespective of the size of transcription unit. Because MS2 signal becomes visible only when elongating RNA polymerase II enters into the 3′ UTR of the gene, measurement of transcription dot allows me to estimate the rate of full-length mRNA production with high temporal resolution. Fluorescent *in situ* hybridization (FISH) assay using MS2 probe showed that authentic patterns of gene expression are maintained in genome editing strains (Figure S1A-C). Importantly, they were all homozygote viable, suggesting that engineering of 3′ UTR does not impede either transcription or translation. This is important for the analysis of developmental genes in early embryos because many of them encode sequence-specific transcription factors and are subject to auto-regulation by their own protein products [21–23].

**Figure 1.**
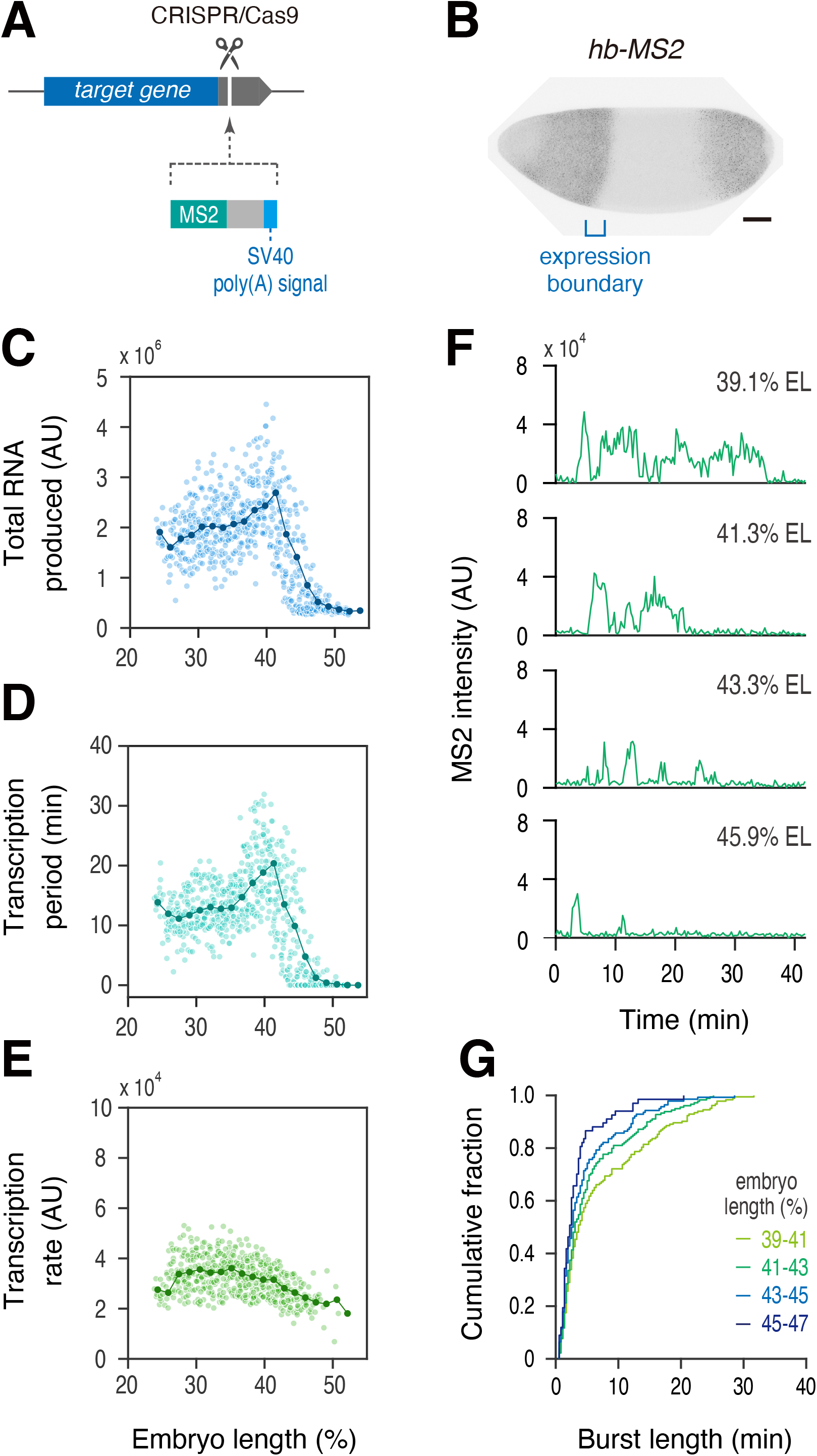
Length of transcription period determines spatial pattern of *hb* expression. (A) Sequence cassette containing 24x MS2 and SV40 poly(A) signal was inserted into the 3′ UTR of target genes by using CRISPR/Cas9. (B) Fluorescent *in situ* hybridization of endogenous *hb-MS2*. Embryo at nc14 was shown. MS2 probe was used for the analysis. Image was cropped and rotated to align embryo (anterior to the left and posterior to the right). Scale bar indicates 50 μm. (C-E) Profiles of total RNA production (C), transcription period (D) and transcription rate (E) as a function of AP position. Line plot indicates mean values in groups of nuclei binned by AP position. In total, 695 nuclei from three independent embryos were analyzed. (F) Representative trajectories of transcription activity of *hb-MS2* in individual nuclei. (G) A cumulative plot showing fraction of bursting events (y axis) and length of transcriptional bursting (x axis). A total of 230 nuclei from three independent embryos was analyzed.

First, expression of endogenous *hb* was visualized with maternally provided MCP-GFP fusion protein from the entry into nuclear cycle 14 (nc14) (Movie S1). Integration of MS2 signal in individual nuclei over time recapitulated the formation of sharp ON/OFF boundary with steep gradient around ~43-47% embryo length (EL) (Figure 1C). To explore the mechanism of pattern formation, quantitative image analysis was performed to determine the total duration of active transcription (transcription period) and the mean MS2 intensity during transcriptional activation (transcription rate) (Figure S1D). I found that the length of transcription period in each nucleus is sufficient to recapitulate spatial patterning of nascent RNA production including the formation of expression boundary (Figure 1D), while the transcription rate was constant regardless of relative location (Figure 1E). To ask how the length of transcription period is regulated, MS2 trajectories in individual nuclei were analyzed. Nuclei located at boundary region were found to produce sporadic transcriptional bursting (Figure 1F; 43.3% EL and 45.9% EL). On the other hand, nuclei near the core expression domain exhibited more continuous activation profiles (Figure 1F; 39.1%EL and 41.3% EL). This trend was clearly seen when analyzed length of every single bursting events in all nuclei at each AP position (Figure 1G), suggesting that nuclei located near the core expression domain produce next burst before clearance of previous bursts to increase the total length of transcription period. Next, I analyzed *hb* expression at the posterior part of embryos (Movie S2). As in anterior expression domain, the length of transcription period highly correlated with the total output, while the transcription rate exhibited only moderate correlation (Figure S2A-C). Individual MS2 trajectories of posterior *hb* exhibited similar transition of bursting profiles along the AP axis (Figure S2D and E). These results are consistent with the idea that the modulation of transcription period is a major determinant of spatial patterning of *hb* expression in early embryos.

Next, I explored transcription dynamics of other gap genes, *gt*, *Kr* and *kni* (Figure S1A). Live imaging analysis revealed that their spatial patterns also arise from the modulation of transcription period as in *hb* locus (Figure 2A and B). Comparison of individual MS2 trajectories revealed differential continuity of bursting activities between boundary and core expression domain (Figure S3). Superimposed profiles of transcription period nicely overlapped with spatial distribution of nascent transcripts within an embryo (Figure 2B), while the transcription rate did not (Figure 2C), again supporting the idea that unique expression patterns of gap genes emerge from the modulation of transcription period. Importantly, each gene was found to exhibit distinct rate of nascent RNA production during transcriptional activation (Figure 2C and G), which appears to contribute to gene-to-gene variability in the total output. For example, posterior *hb* exhibits longer transcription period than *kni* (Figure 2B), but does not produce proportionally higher level of output since the transcription rate is nearly twice smaller (Figure 2C and G). I therefore suggest that the length of transcription period defines basic spatial patterning of gene expression and the transcription rate contributes to differential output among gap genes.

**Figure 2.**
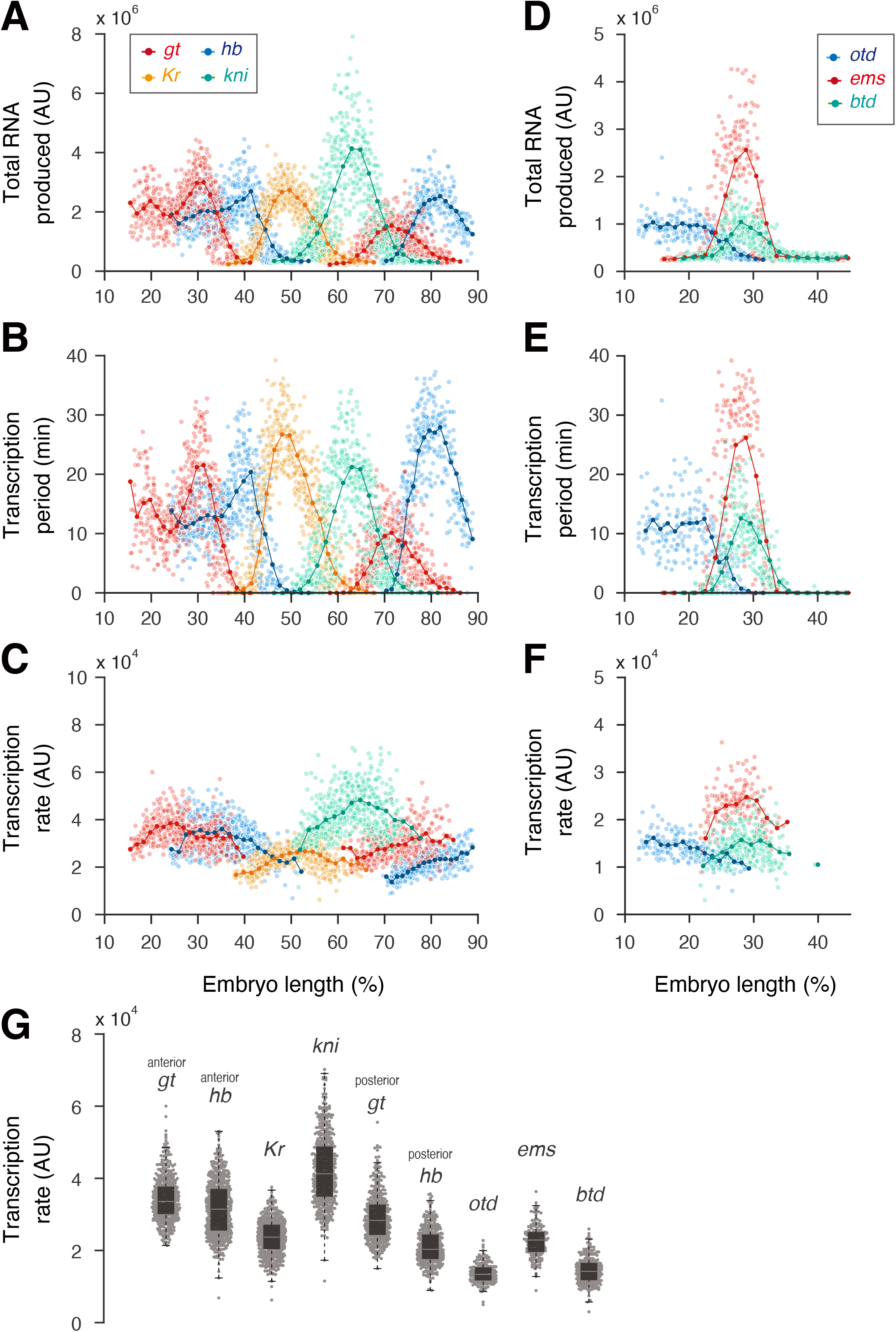
Spatial patterns of gap genes arise from the modulation of transcription period. (A-C) Imposed profiles of total RNA production (A), transcription period (B) and transcription rate (C) as a function of AP position. Line plots indicate mean values in groups of nuclei binned by AP position. A total of 546, 695, 759, 740, 608, 429 nuclei, respectively, were analyzed from three independent embryos for anterior *gt*, anterior *hb*, *Kr*, *kni*, posterior *gt* and posterior *hb*. Plots of *hb* are same as the plots in Figure 1C-E and Figure S2A-C. (D-F) Imposed profiles of total RNA production (D), transcription period (E) and transcription rate (F) as a function of AP position. Line plots indicate mean values in groups of nuclei binned by AP position. A total of 270, 481, 668 nuclei, respectively, were analyzed from three independent embryos for *otd*, *ems* and *btd*. (G) Boxplot showing the distribution of transcription rate shown in (C) and (F). The box indicates the lower (25%) and upper (75%) quantiles, and the solid line indicates the median. Whiskers extend to the most extreme, non-outlier data points.

To further explore regulatory mechanism of other key AP patterning genes, transcription dynamics of endogenous head gap genes, *orthodenticle* (*otd*), *buttonhead* (*btd*) and *empty spiracles* (*ems*) was visualized (Figure S1B). Live imaging analysis revealed that spatial patterns of head gap genes also arise from the modulation of transcription period rather than the transcription rate (Figure 2D-F). Same conclusion was further supported by the analysis of endogenous pair-rule genes, *even-skipped* (*eve*), *fushi-tarazu* (*ftz*), *hairy* (*h*), *paired* (*prd*) and *runt* (*run*) (Figure S1C and S4A). Comparison of transcription rates revealed that there are ~3.1-fold and ~2.7-fold variations among gap and pair-rule genes (Figure 2G and Figure S4B). Among these, rate of *hb* transcription was found to largely differ between anterior and posterior expression domains (Figure 2C and G). In contrast, rate of *gt* transcription was similar between anterior and posterior regions (Figure 2C and G). Transcription of *gt* is driven from the common promoter regardless of the expression domain. On the other hand, anterior expression of *hb* is driven from the proximal P2 promoter [24], while the posterior expression is also driven from the distal P1 promoter (Figure 3A) [25, 26], suggesting that differential promoter usage within a same gene mediates switching of transcription rate in response to different transcriptional activators present along the AP axis.

**Figure 3.**
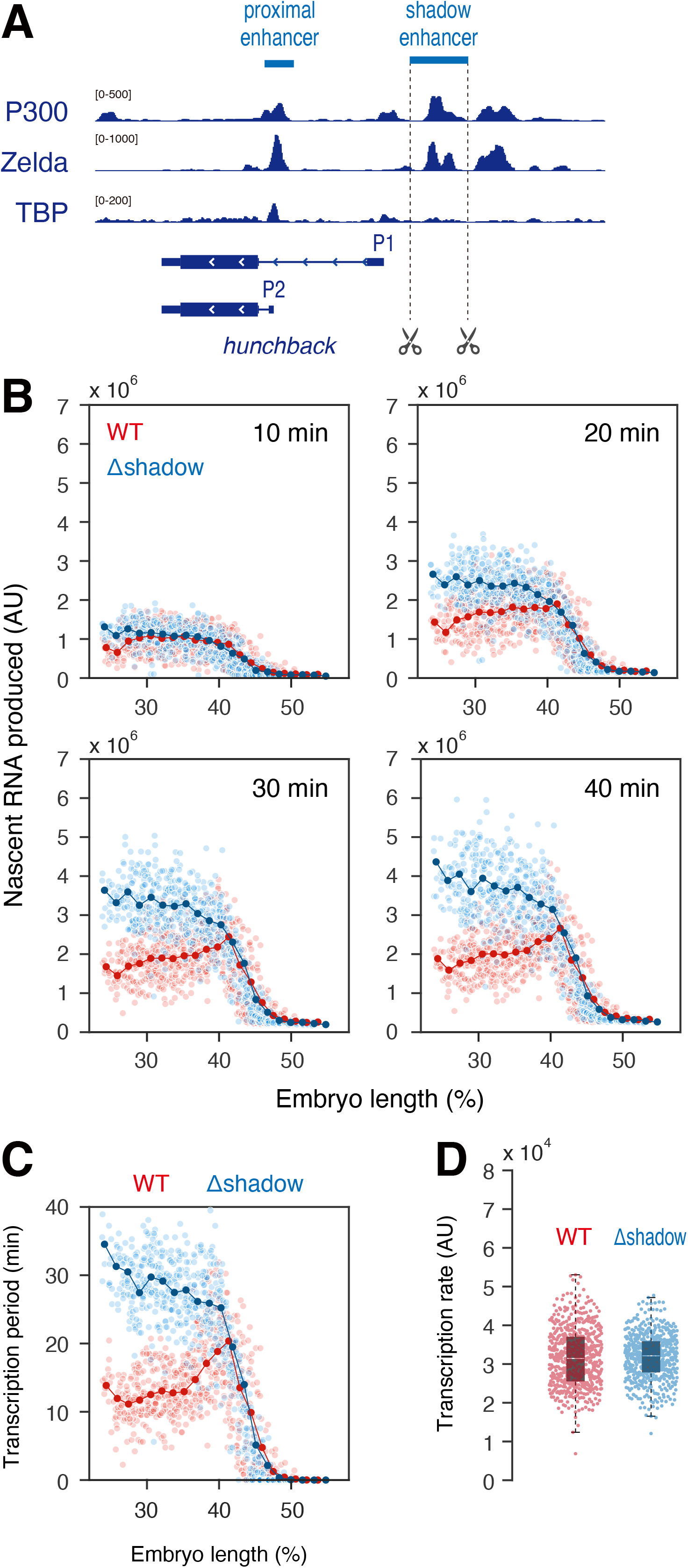
Enhancer competition at the *hb* locus. (A) Organization of the endogenous *hb* locus. P300 ChIP-seq data from nc12 to nc14 WT embryos [50], Zelda ChIP-seq data from 2- to 4-h WT embryos [51], and TBP ChIP-seq data from 2- to 4-h WT embryos [52] were visualized with Integrative Genomics Viewer (IGV). (B) Time-course measurement of nascent *hb* RNA production in WT and deletion mutant as a function of AP position. In total, 695 and 681 nuclei from three independent embryos were analyzed. (C) Profile of transcription period in WT and deletion mutant. Line plot indicates mean values in groups of nuclei binned by AP position. In total, 695 and 681 nuclei from three independent embryos were analyzed. Plot of WT is same as the plot in Figure 1D. (D) Boxplot showing the distribution of transcription rates in WT and deletion mutant. The box indicates the lower (25%) and upper (75%) quantiles, and the solid line indicates the median. Whiskers extend to the most extreme, non-outlier data points. In total, 695 and 681 nuclei from three independent embryos were analyzed.

Gap genes are typically regulated by two redundant enhancers that can produce overlapping patterns in early embryos [27]. While previous transgene assay suggested that redundant enhancers foster robustness of gene expression [28, 29] or help to increase transcriptional output [30], it remains unclear if they can simultaneously interact with target promoter. Anterior expression of *hb* is controlled not only by classical proximal enhancer [24, 31], but also by newly-identified distal redundant enhancer, or shadow enhancer [27] (Figure 3A). To visualize the role of shadow enhancer at the endogenous locus, second round of genome editing was performed using *hb-MS2* strain to remove corresponding regulatory sequence (Figure 3A). Unexpectedly, live imaging analysis of resulting mutant embryos revealed that *hb* starts to produce exceptionally high level of nascent transcripts upon deletion of shadow enhancer (Figure 3B and Movie S3). Time-course measurement revealed that this ectopic activation takes place ~10-20 min after entry into nc14 (Figure 3B). Importantly, removal of shadow enhancer increased the length of transcription period without changing the transcription rate (Figure 3C and D). These results are consistent with the idea that distal shadow enhancer interferes with gene activation by proximal enhancer to downregulate *hb* expression at internal region of expression domain. It has been shown that spatial limit of *hb* expression is determined by the gradient of maternal Bicoid activator [24]. I found that the level of RNA production at boundary region (~43-47% EL) was unaffected by the removal of shadow enhancer (Figure 3B), suggesting that proximal enhancer alone is sufficient for responding to diminishing level of Bicoid, and that shadow enhancer does not impede this activity.

Lastly, I analyzed time-dependent shifting of gap gene expression during nc14. Consistent with previous studies using fixed embryos [7], live imaging data captured anterior shifting of transcription activities of *Kr*, *kni*, posterior *gt* and *hb* (Figure 4A). However, simple accumulation of nascent transcripts over time was found to mask these shifting effects (Figure 4B), which is incompatible with previous quantitative FISH analysis [7, 32]. Early *Drosophila* embryo is characterized by rapid rates of mRNA degradation. Half-life of newly synthesized mRNAs in early nc14 embryos is estimated to be ~13-14 min [33, 34]. Using live imaging data, I simulated effects of mRNA degradation by considering rapid rate of mRNA decay (t_1/2_ = 13 min). Simulated data augmented anterior shifting of expression domains (Figure 4C), suggesting that both *de novo* RNA synthesis and degradation rates contribute to dynamic shifting of positional information encoded by the gap genes. Simulated data also suggest that rapid mRNA degradation helps refinement of initial broad expression domain of anterior *gt* and *hb* into narrower stripe patterns (Figure 4C). Importantly, *hb* mutant lacking distal shadow enhancer can only produce static pattern of gene expression even in the presence of rapid mRNA decay (Figure 4D and E). I therefore suggest that transcriptional interference by distal shadow enhancer facilitates dynamic refinement of expression domain in early embryos.

**Figure 4.**
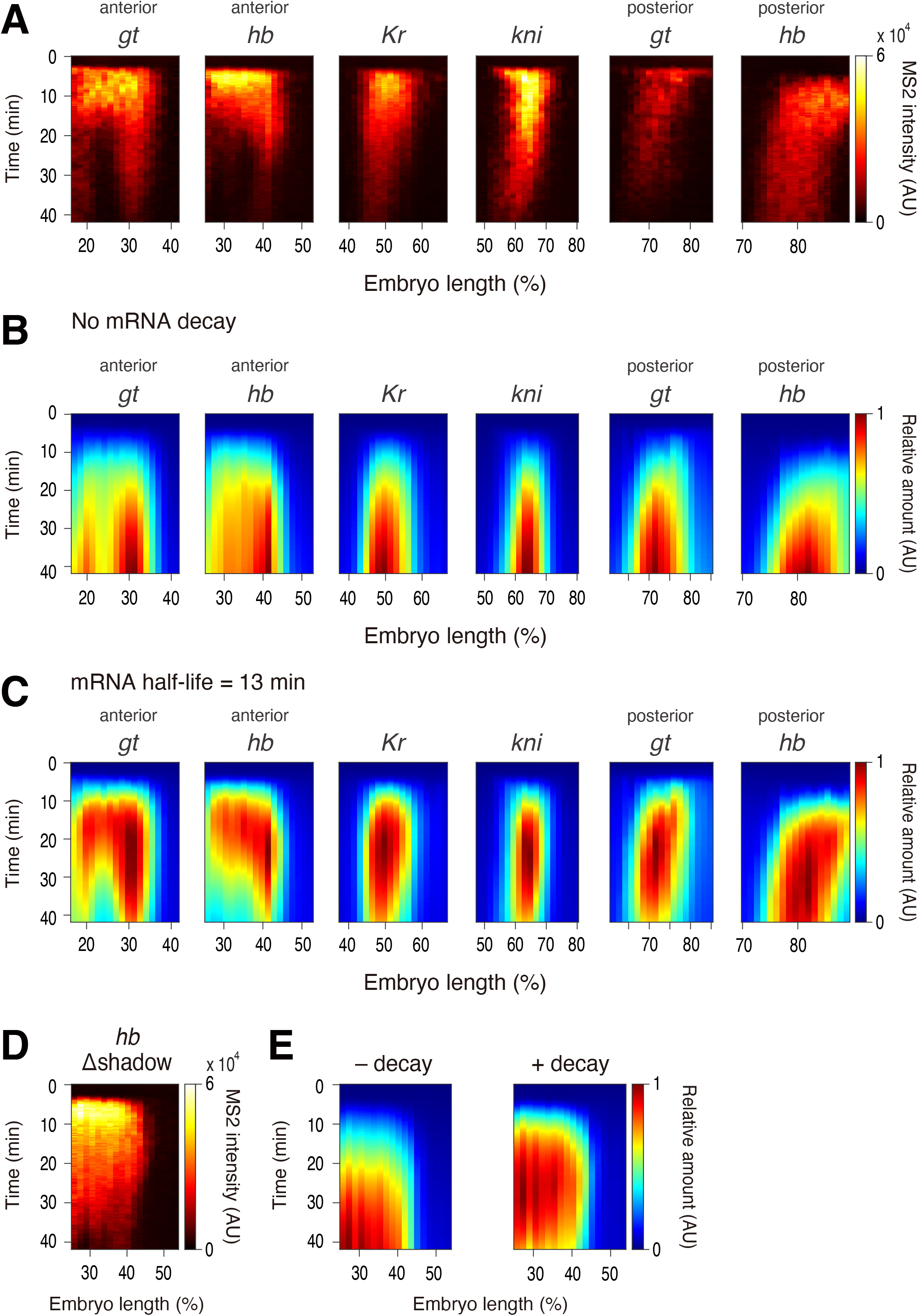
Dynamic refinement of gap gene expression. (A) Averaged MS2 signal in groups of nuclei binned by AP position over space and time. Data set shown in Figure 2 was used for the analysis. (B-C) Averaged cumulative RNA production in groups of nuclei binned by AP position in the absence (B) or presence of mRNA decay (C). Data set shown in Figure 2 was used for the analysis. (D) Averaged MS2 signal in groups of nuclei binned by AP position. Data set shown in Figure 3 was used for the analysis. (E) Averaged cumulative RNA production in groups of nuclei binned by AP position in the absence (left) or presence of mRNA decay (right). Data set shown in Figure 3 was used for the analysis.

In this study, I provided evidence that the modulation of transcription period is a major determinant of spatial patterning of gene expression in *Drosophila* embryos. The length of transcription period is regulated by the continuity of bursting activities in individual nuclei, with core expression domain producing more bursts than boundary region to increase total duration of active transcription (Figure 1, Figure S2 and S3). Consistent with previous results [35, 36], enhancers appear to be responsible for successive induction of transcriptional bursting because removal of *hb* shadow enhancer specifically altered the length of transcription period without changing the transcription rate (Figure 3C and D). Previous transgene assay suggested that *hb* shadow enhancer is ~2-fold weaker than proximal enhancer [30]. Genome-editing analysis in this study revealed that the deletion of shadow enhancer causes ~2-fold increase in the level of total RNA production (Figure 3B), suggesting that interaction between P2 promoter and strong proximal enhancer is competitively blocked by distal weak enhancer to downregulate *de novo* RNA synthesis and facilitate refinement of initial expression domain during nc14. Antagonistic relationship of *hb* enhancers suggests that they do not form common “transcription hubs” for gene activation, which differs from cooperative action of super enhancers in mammalian cells [37–39]. Regulation of *hb* is also unique from enhancer competition mechanism seen at other locus [30, 40] in a point that distal weak enhancer sequesters interaction with target promoter from remote location despite its “topological disadvantage” over strong proximal enhancer. It might be possible that transcriptional interference by *hb* shadow enhancer helps subsequent recruitment of distal stripe enhancer to the promoter region at late nc14 [41]. Supporting this view, clear stripe pattern was not observed in the deletion mutant (Figure 4D and E, Movie S3).

This study also showed that the transcription rate is relatively constant across expression domain for all tested genes, which is consistent with previous observation that transcriptional activators and repressors mainly regulate the frequency, but not the amplitude, of transcriptional bursting [36, 42]. When compared different genes, each of them exhibits distinct rate of transcription (Figure 2G and Figure S4B). This appears to contribute to gene-to-gene variability in the level of total RNA production (Figure 2A-F and Figure S4A). Transcription rates might be modulated by core promoter sequences because different usage of *hb* promoter largely altered its activity within an embryo (Figure 2C and G). Recent single molecule FISH and theoretical modeling approach suggested that transcription initiation rates are constant among four major gap genes in nc13 embryos [43]. It can be possible that transcription rates of gap genes in earlier nuclear cycle are equally “boosted-up” to enable efficient production of nascent transcripts within a short duration of interphase (~10-12 min). I suggest that differential rates of transcription in nc14 help to enrich regulatory information encoded by the gap and pair-rules genes for specification of individual body segments in developing embryos.

## Acknowledgment

I thank Manuel Cambón for developing image analysis code, Hitomi Takishita for help with screening of genome-editing strains and fly husbandry, and Misako Sato for fly husbandry. I am also grateful to members of the Fukaya laboratory for discussion. This study was funded by the Grant-in-Aid for Scientific Research on Innovative Areas (Research in a Proposed Research Area) (20H05357), the Grants-in-Aid for Scientific Research (B) (19H03154), the Grants-in-Aid for Challenging Research (Exploratory) (19K22378) from the Japan Society for the Promotion of Science, the Grants-in-Aid for Leading Initiative for Excellent Young Researchers from the Ministry of Education, Culture, Sports, Science and Technology in Japan, Tomizawa Jun-ichi & Keiko Fund of Molecular Biology Society of Japan for Young Scientist, research grants from the Mochida Memorial Foundation for Medical and Pharmaceutical Research, the Nakajima Foundation, the Inamori Foundation, the Takeda Science Foundation and the Sumitomo Foundation.

## Author Contributions

T.F designed and performed the experiments, analyzed the data and wrote the manuscript.

## Contact for reagent and resource sharing

Further information and requests for resources and reagents should be directed to and will be fulfilled by the Lead Contact, Takashi Fukaya.

## Experimental model and subject details

In all imaging experiments, I studied *Drosophila melanogaster* embryos at nuclear cycle 14. The following fly lines were used in this study: *nos>MCP-GFP, His2Av-mRFP/CyO* [44], *hb-MS2* (this study), *gt-MS2* (this study), *Kr-MS2* (this study), *kni-MS2* (this study), *otd-MS2* (this study), *ems-MS2* (this study), *btd-MS2* (this study), *h-MS2* (this study), *run-MS2* (this study), *prd-MS2* (this study), *hb-MS2 Δshadow enhancer* (this study), *ftz-MS2* [45], and *eve-MS2* [45].

## Method Detail

### MS2-tagging of endogenous locus by CRISPR/Cas9

pCFD3 gRNA expression plasmid and pBS-MS2-dsRed donor plasmid were co-injected using *nanos-Cas9* strains [46]. Microinjection was performed as previously described [47]. In brief, 0-1 h embryos were collected and dechorionated with bleach. Aligned embryos were dried with silica gel for ~7 min and covered with FL-100-1000CS silicone oil (Shin-Etsu Silicone). Subsequently, microinjection was performed using FemtoJet 5247 (Eppendorf) and DM IL LED inverted microscope (Leica) equipped with M-152 Micromanipulator (Narishige). Injection mixture typically contains 500 ng/μl pCFD3 gRNA expression plasmid, 500 ng/μl pBS-MS2-dsRed donor plasmid, 5 mM KCl, 0.1 mM phosphate buffer, pH 6.8. 3xP3-dsRed marker was used for screening.

### Enhancer deletion by CRISPR/Cas9

pCFD3 gRNA expression plasmids, pBS-3xP3-GFP donor plasmid and pBS-hsp70-Cas9 plasmid (addgene #46294) were co-injected to homozygote *hb-MS2* embryos. Microinjection was performed as described in previous section. Injection mixture contains 500 ng/μl pCFD3 gRNA expression plasmids, 500 ng/μl pBS-3xP3-GFP donor plasmid, 500 ng/μl pBS-hsp70-Cas9 plasmid, 5 mM KCl, 0.1 mM phosphate buffer, pH 6.8. 3xP3-GFP marker was used for subsequent screening. Deletion has been confirmed by PCR analysis of genomic DNA purified from the resulting mutant.

### Preparation of MS2 probe for *in situ* hybridization

Antisense RNA probe labeled with digoxigenin (DIG RNA Labeling Mix 10 × conc, Roche) was *in vitro* transcribed using T3 RNA polymerase (NEB). Templated DNA for MS2 probe was prepared by linearizing pBlueScript-MS2 plasmid [48] with EcoRI.

### Fluorescence *in situ* hybridization

Embryos were dechorionated and fixed in fixation buffer (1 ml of 5x PBS, 4 ml of 37% formaldehyde and 5 ml of Heptane) for ~25 min at room temperature. Antisense RNA probe labeled with digoxigenin was used. Hybridization was performed at 55 °C overnight in hybridization buffer (50% formamide, 5x SSC, 50 μg/ml Heparin, 100 μg/ml salmon sperm DNA, 0.1% Tween-20). Subsequently, embryos were washed with hybridization buffer at 55 °C and incubated with Western Blocking Reagent (Roche) at room temperature for ~2 hours. Then, embryos were incubated with sheep anti-digoxigenin (Roche) at 4 °C for overnight, followed by incubation with Alexa Fluor 555 donkey anti-sheep (Invitrogen) fluorescent secondary antibody at room temperature for ~2 hour. Embryos were mounted in ProLong Gold Antifade Mountant (Thermo Fisher Scientific). Imaging was performed on a Zeiss LSM 900 confocal microscope. Plan-Apochromat 20x / 0.8 N.A. objective was used. Images were captured in 16-bit. Maximum projections were obtained for all z-sections, and resulting images were shown. Brightness of images was linearly adjusted using Fiji (https://fiji.sc).

### MS2 live imaging

*MCP-GFP, His2Av-mRFP/CyO* virgin females were mated with males carrying the MS2 allele. The resulting embryos were dechorionated and mounted between a polyethylene membrane (Ube Film) and a coverslip (18 mm × 18 mm), and embedded in FL-100-450CS (Shin-Etsu Silicone). Embryos were imaged using a Zeiss LSM 900. Room temperature was kept in between 22.0 to 23.0 °C during imaging. Plan-Apochromat 40x / 1.4 N.A. oil immersion objective was used. At each time point, a stack of 26 images separated by 0.5 μm was acquired. Typical time resolution of resulting maximum projection was 0.28 min. In subsequent image analysis, all movies were considered to have same time resolution (0.28 min/frame). Images were captured in 16-bit. Images were typically taken from the end of nc13 to the onset of gastrulation at nc14. During imaging, data acquisition was occasionally stopped for a few seconds to correct z-position, and data were concatenated afterwards. For each cross, three independent embryos were analyzed. The same laser power and microscope setting were used throughout the study.

### Plasmids

#### pCFD3-dU6-hb 3′UTR

Two DNA oligos (5′-GTC GAT AGG AGA CAG ATT GAA AGG-3′) and (5′-AAA CCC TTT CAA TCT GTC TCC TAT-3′) were annealed and inserted into the pCFD3-dU6:3gRNA vector (addgene # 49410) using BbsI sites.

#### pBS-hb 5′Arm-24xMS2-dsRed-SV40-hb 3′Arm

A DNA fragment containing 3′ homology arm of *hb* was amplified from genomic DNA using primers (5′-TTT AAT CTA GAT TCA ATC TGT CTC CTA TAC ACT C-3′) and (5′-TTA AAG CGG CCG CCT TGG GTA TTT AGC ACA TGA TGG AG-3′), and digested with XbaI and NotI. The resulting fragment was inserted into pBS-MS2-dsRed [49]. Subsequently, a DNA fragment containing 5′ homology arm of *hb* was amplified from genomic DNA using primers (5′-TTT AAG GTA CCC TCC AGA TGC TGG CCG CCC AAC-3′) and (5′-TTA AAG TCG ACA GGT GGC TAC AGT TCA ATA CAG TTA-3′), and digested with KpnI and SalI. The resulting fragment was inserted into the plasmid.

#### pCFD3-dU6-gt 3′UTR

Two DNA oligos (5′-GTC GCT CTT GAT GCG TTG AAC GCG-3′) and (5′-AAA CCG CGT TCA ACG CAT CAA GAG-3′) were annealed and inserted into the pCFD3-dU6:3gRNA vector (addgene # 49410) using BbsI sites.

#### pBS-gt 5′Arm-24xMS2-dsRed-SV40-gt 3′Arm

A DNA fragment containing 5′ homology arm of *gt* was amplified from genomic DNA using primers (5′-TTT AAC TCG AGG ACT CCC TCG CCG TAT CAA ACA AGC-3′) and (5′-TTA AAG AAT TCG TTC AAC GCA TCA AGA GAG GAG TGG-3′), and digested with XhoI and EcoRI. The resulting fragment was inserted into pBS-MS2-dsRed [49]. Subsequently, a DNA fragment containing 3′ homology arm of *gt* was amplified from genomic DNA using primers (5′-TTT AAA CTA GTG CGA GGA TCC GAA TCG TAT GTC CAG-3′) and (5′-TTA AAC CGC GGC GAC CCT ATT TTC CTC CGA CGT CT-3′), and digested with SpeI and SacII. The resulting fragment was inserted between the the plasmid.

#### pCFD3-dU6-Kr 3′UTR

Two DNA oligos (5′-GTC GAT TAG GGC TAT GTA CAA TT-3′) and (5′-AAA CAA TTG TAC ATA GCC CTA AT-3′) were annealed and inserted into the pCFD3-dU6:3gRNA vector (addgene # 49410) using BbsI sites.

#### pBS-Kr 5′Arm-24xMS2-dsRed-SV40-Kr 3′Arm

A DNA fragment containing 5′ homology arm of *Kr* was amplified from genomic DNA using primers (5′-TTT AAC TCG AGC AGA CCG AGA TCA GCA TGA GTG TA-3′) and (5′-TTA AAG AAT TCA TTC GGT GTG GTA CTG GCC TAA TG-3′), and digested with XhoI and EcoRI. The resulting fragment was inserted into pBS-MS2-dsRed [49]. Subsequently, a DNA fragment containing 3′ homology arm of *Kr* was amplified from genomic DNA using primers (5′-TTT AAA CTA GTT GTA CAT AGC CCT AAT CAG TTT TC-3′) and (5′-TTA AAG CGG CCG CCG ATA CAG AGG TAT TGG AGG TAT-3′), and digested with SpeI and NotI. The resulting fragment was inserted into the plasmid.

#### pCFD3-dU6-otd 3′UTR

Two DNA oligos (5′-GTC GTC CTT TGC GAT CGT ATT TGT-3′) and (5′-AAA CAC AAA TAC GAT CGC AAA GGA-3′) were annealed and inserted into the pCFD3-dU6:3gRNA vector (addgene # 49410) using BbsI sites.

#### pBS-otd 5′Arm-24xMS2-dsRed-SV40-otd 3′Arm

A DNA fragment containing 5′ homology arm of *otd* was amplified from genomic DNA using primers (5′-TTT AAC TCG AGG GAT GCC GGT GGT GAT ATT GGT GCC-3′) and (5′-TTA AAG AAT TCA ATA CGA TCG CAA AGG AAG TGT ATC-3′), and digested with XhoI and EcoRI. The resulting fragment was inserted into pBS-MS2-dsRed [49]. Subsequently, a DNA fragment containing 3′ homology arm of *otd* was amplified from genomic DNA using primers (5′-TTT AAA CTA GTT GTT GGA AAT AGG AAA TTC AAG TTC-3′) and (5′-TTA AAC CGC GGG CTG ATT TGG GTT TGT ATT CTC-3′), and digested with SpeI and SacII. The resulting fragment was inserted into the plasmid.

#### pCFD3-dU6-ems 3′UTR

Two DNA oligos (5′-GTC GCG GCC AGG AGA TAG TCC TG-3′) and (5′-AAA CCA GGA CTA TCT CCT GGC CG-3′) were annealed and inserted into the pCFD3-dU6:3gRNA vector (addgene # 49410) using BbsI sites.

#### pBS-ems 5′Arm-24xMS2-dsRed-SV40-ems 3′Arm

A DNA fragment containing 5′ homology arm of *ems* was amplified from genomic DNA using primers (5′-TTT AAC TCG AGC GCC CCT TTC CCA TGG GAC CCG GA-3′) and (5′-TTA AAG AAT TCG ACT ATC TCC TGG CCG CTT CTC TC-3′), and digested with XhoI and EcoRI. The resulting fragment was inserted into pBS-MS2-dsRed [49]. Subsequently, a DNA fragment containing 3′ homology arm of *ems* was amplified from genomic DNA using primers (5′-TTT AAG CTA GCC TGC GGC GGT CCG TGA GTT CCT T-3′) and (5′-TTA AAC CGC GGC CCG CTC TTT TCA GTC GCG GCC GCA-3′), and digested with NheI and SacII. The resulting fragment was inserted into the plasmid.

#### pCFD3-dU6-btd 3′UTR

Two DNA oligos (5′-GTC GTT CCC CCA ATC TAG TAG CTA-3′) and (5′-AAA CTA GCT ACT AGA TTG GGG GAA-3′) were annealed and inserted into the pCFD3-dU6:3gRNA vector (addgene # 49410) using BbsI sites.

#### pBS-btd 5′Arm-24xMS2-dsRed-SV40-btd 3′Arm

A DNA fragment containing 5′ homology arm of *btd* was amplified from genomic DNA using primers (5′-TTT AAG GTA CCC GCG CAG CGA TCA CCT CAG CAA GC-3′) and (5′-TTA AAC TCG AGC TAC TAG ATT GGG GGA AAA AAA-3′), and digested with KpnI and XhoI. The resulting fragment was inserted into pBS-MS2-dsRed [49]. Subsequently, a DNA fragment containing 3′ homology arm of *btd* was amplified from genomic DNA using primers (5′-TTT AAA CTA GTC TAA GGC ATT TTT TTT TGC ATG TC-3′) and (5′-TTA AAC CGC GGC CCG TTT AAT TAT AAA CTC TGT G-3′), and digested with SpeI and SacII. The resulting fragment was inserted into the the plasmid.

#### pCFD3-dU6-h 3′UTR

Two DNA oligos (5′-GTC GAG ACA TTT CAC ATC ATT CGC-3′) and (5′-AAA CGC GAA TGA TGT GAA ATG TCT-3′) were annealed and inserted into the pCFD3-dU6:3gRNA vector (addgene # 49410) using BbsI sites.

#### pBS-h 5′Arm-24xMS2-dsRed-SV40-h 3′Arm

A DNA fragment containing 3′ homology arm of *h* was amplified from genomic DNA using primers (5′-TTT AAT CTA GAC GCC GGG ATT GCG CAA ATG TTG CT-3′) and (5′-TTA AAG CGG CCG CGG CCT ATC GAA CGG ATG TGT GAA-3′), and digested with XbaI and NotI. The resulting fragment was inserted into pBS-MS2-dsRed [49]. Subsequently, a DNA fragment containing 5′ homology arm of *h* was amplified from genomic DNA using primers (5′-TTT AAG GTA CCG CCG GCT CGC CAC TCC AAA TTG G-3′) and (5′-TTA AAC TCG AGA ATG ATG TGA AAT GTC TAC GCG C-3′), and digested with KpnI and XhoI. The resulting fragment was inserted into the plasmid.

#### pCFD3-dU6-run 3′UTR

Two DNA oligos (5′-GTC GCG GCG ACG CAG AGC GGC AAG-3′) and (5′-AAA CCT TGC CGC TCT GCG TCG CCG-3′) were annealed and inserted into the pCFD3-dU6:3gRNA vector (addgene # 49410) using BbsI sites.

#### pBS-run 5′Arm-24xMS2-dsRed-SV40-run 3′Arm

A DNA fragment containing 5′ homology arm of *run* was amplified from genomic DNA using primers (5′-TTT AAC TCG AGG CCG CCA ACC AAA TCC CCC ACC AT-3′) and (5′-TTA AAG AAT TCG CCG CTC TGC GTC GCC GCT GTT ATA-3′), and digested with XhoI and EcoRI. The resulting fragment was inserted into pBS-MS2-dsRed [49]. Subsequently, a DNA fragment containing 3′ homology arm of *run* was amplified from genomic DNA using primers (5′-TTT AAT CTA GAA AGC GGG CAA AAA TTA TAA TTG TG-3′) and (5′-TTA AAG CGG CCG CGC TAA TGC TCG GAG ATA AGT TAA-3′), and digested with XbaI and NotI. The resulting fragment was inserted into the plasmid.

#### pCFD3-dU6-prd 3′UTR

Two DNA oligos (5′-GTC GAT CCA CCA CCT ACT CCT CC-3′) and (5′-AAA CGG AGG AGT AGG TGG TGG AT-3′) were annealed and inserted into the pCFD3-dU6:3gRNA vector (addgene # 49410) using BbsI sites.

#### pBS-prd 5′Arm-24xMS2-dsRed-SV40-prd 3′Arm

A DNA fragment containing 5′ homology arm of *prd* was amplified from genomic DNA using primers (5′-TTT AAC TCG AGC GGC GTG CTC GTC TCC GCA AGC AGC-3′) and (5′-TTA AAG AAT TCG GAG TAG GTG GTG GAT CCG TGT CCC-3′), and digested with XhoI and EcoRI. The resulting fragment was inserted into pBS-MS2-dsRed [49]. Subsequently, a DNA fragment containing 3′ homology arm of *prd* was amplified from genomic DNA using primers (5′-TTT AAA CTA GTT CCA GGA GCA GGA GCA GGT GTC ACC-3′) and (5′-TTA AAG CGG CCG CGT GCA ATC CGT GCA CAG ATC TTT GC-3′), and digested with SpeI and NotI. The resulting fragment was inserted into the plasmid.

#### pCFD3-dU6-hb shadow enhancer-1

Two DNA oligos (5′-GTC GTA TCA TTG TTA GCT TGA CA-3′) and (5′-AAA CTG TCA AGC TAA CAA TGA TA-3′) were annealed and inserted into the pCFD3-dU6:3gRNA vector (addgene # 49410) using BbsI sites.

#### pCFD3-dU6-hb shadow enhancer-2

Two DNA oligos (5′-GTC GCA ATT AGA AAC CTA TCC AAA-3′) and (5′-AAA CTT TGG ATA GGT TTC TAA TTG-3′) were annealed and inserted into the pCFD3-dU6:3gRNA vector (addgene # 49410) using BbsI sites.

#### pBS-hb shadow enhancer 5′Arm-attP-dsRed-SV40-hb shadow enhance 3′Arm

A DNA fragment containing 5′ homology arm of *hb* shadow enhancer was amplified from genomic DNA using primers (5′-TTT AAC TCG AGG CAC AAG CAC GAA GTT TAA TTG-3′) and (5′-TTA AAG AAT TCC AAG CTA ACA ATG ATA CAT TTT CCG-3′), and digested with XhoI and EcoRI. The resulting fragment was inserted into pBS-3xP3-GFP [45]. Subsequently, a DNA fragment containing 3′ homology arm of *hb* shadow enhancer was amplified from genomic DNA using primers (5′-TTT AAA CTA GTA AAA GGA TAG GTT CAA TGA TGT TAG-3′) and (5′-TTA AAG CGG CCG CGC CAG CTA CCT GCC CGC ACC GTT GG-3′), and digested with SpeI and NotI. The resulting fragment was inserted into the plasmid.

### Image analysis

All the image processing methods and analysis were implemented in MATLAB (R2019b, MathWorks).

### Segmentation of nuclei

For each time point, maximum projections were obtained for all 26 z-sections per image. His2Av-mRFP was used to segment nuclei. His2Av images were first blurred with Gaussian filtering to generate smoothed images. Pixels expressing intensity higher than 5% of the global maxima in the histogram of His2Av channel were removed. Processed images were converted into binary images using a custom threshold-adaptative segmentation algorithm. Threshold values were determined at each time frame by taking account of 1) histogram distribution of His2Av channel and 2) the number and size of resulting connected components. Boundaries of components were then modified to locate MS2 transcription dots inside of nearest nuclei. In brief, pixels with intensity twice larger than mean intensity in MS2 channel were considered as active transcription dots, and new binary images were created for each time frame. The Euclidean distances between the centroid of binarized transcription dot and all boundaries of segmented nuclei were calculated. Boundary of the nucleus with the smallest Euclidean distance was modified in order to capture transcription dot within a nucleus. Subsequently, centroids of connected components in nuclei segmentation channel were used to compute the Voronoi cells of the image. Resulting binary images were manually corrected by using Fiji (https://fiji.sc).

### Tracking of nuclei

Nuclei tracking was done by finding the object with minimal movement across the frames of interest. For each nucleus in a given frame, the Euclidean distances between the centroids of the nucleus in current time frame and the nuclei in previous time frame were determined. The nucleus with the minimum Euclidean distance was considered as a same lineage.

### Recording of MS2 signal

Maximum projections of raw images were used to record MS2 fluorescence intensities. Using segmented regions, fluorescence intensities within each nucleus were extracted. Signals of MS2 transcription dots were determined by taking integral in a 3 × 3 pixel region centering the brightest pixel within a nucleus after subtracting median fluorescence intensity within a nucleus as a background. Subsequently, minimum MS2 intensities were determined for individual trajectories and subtracted to make the baseline zero.

### Quantification of output, transcription period and transcription rate

For each time frame, nuclei with MS2 intensity above the threshold were considered as active using trajectories after smoothing. Threshold was determined for each gene by calculating the 15% of the maximum intensity across all smoothed trajectories from three independent embryos. Length of transcription period was determined by calculating total active duration for each nucleus. Transcription rate was determined as a mean MS2 intensity during active state using raw trajectories. Total RNA production was measured by taking the area under the raw trajectory. Nuclei were binned into 20 groups by their relative AP position, and mean values were determined for each parameter.

### Description of burst length

For each time frame, nuclei with MS2 intensity above the threshold were considered to be bursting using trajectories after smoothing. Threshold was determined for each gene by calculating the 10% of the maximum MS2 intensity across all smoothed trajectories from three independent embryos. From each trajectory, duration of every single bursting events was measured. Bursts that persist only for a single timeframe were considered as a noise and excluded from the analysis.

### Heatmap analysis of instantaneous MS2 intensity and cumulative RNA production

For each time point, mean MS2 intensities among AP-binned nuclei were determined using raw trajectories and shown as a heatmap. Newly synthesized mRNAs were considered to be linearly degraded with a half-life of 13 min according to previous experimental measurements [33, 34]. Amount of undegraded mRNAs at each time point was calculated using raw MS2 trajectories. For each time point, mean cumulative RNA levels among AP-binned nuclei were determined and shown as a heatmap.

### False-coloring by instantaneous MS2 signals

MS2 signal intensity at each nucleus was measured as described in the previous section. Using segmentation mask, individual nuclei were false-colored with the pixel intensity proportional to the instantaneous MS2 signal at given time in a given nucleus. Resulting image was then colored and layered over the maximum projected image of His2Av-mRFP.

## Supplemental Figure Legends

**Figure S1.**
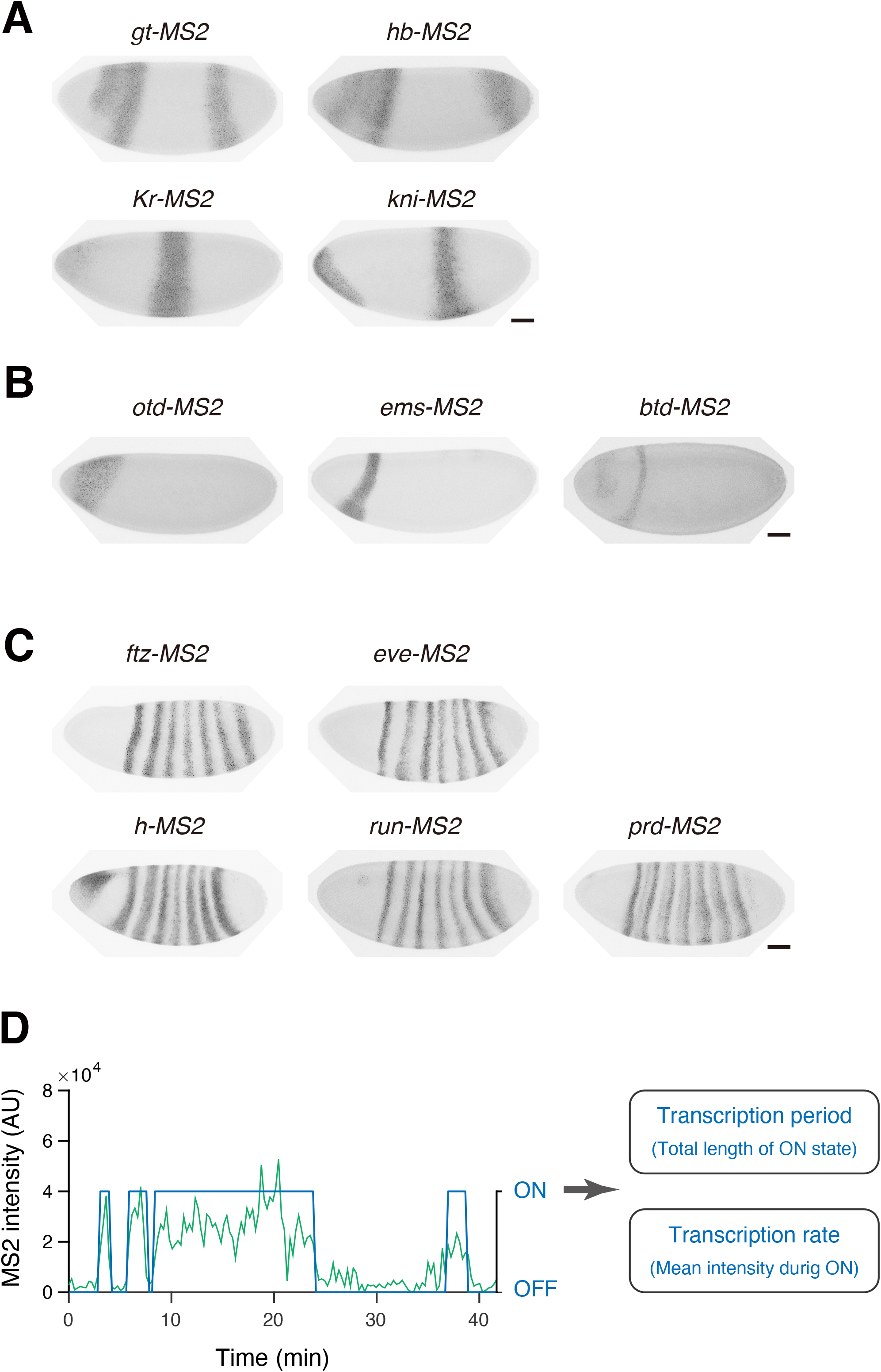
Distribution of MS2-tagged transcripts in genome editing strains. (A) Fluorescent *in situ* hybridization of endogenous *gt-MS2*, *hb-MS2*, *Kr-MS2* and *kni-MS2*. Embryo at nc14 was shown. MS2 probe was used for the analysis. Image was cropped and rotated to align embryos (anterior to the left and posterior to the right). Scale bar indicates 50 μm. (B) Fluorescent *in situ* hybridization of endogenous *otd-MS2*, *ems-MS2* and *btd-MS2*. Embryo at nc14 was shown. MS2 probe was used for the analysis. Image was cropped and rotated to align embryos (anterior to the left and posterior to the right). Scale bar indicates 50 μm. (C) Fluorescent *in situ* hybridization of endogenous *ftz-MS2*, *eve-MS2*, *h-MS2*, *run-MS2* and *prd-MS2*. Embryo at nc14 was shown. MS2 probe was used for the analysis. Image was cropped and rotated to align embryos (anterior to the left and posterior to the right). Scale bar indicates 50 μm. (D) Transcription period and transcription rate were determined for individual nuclei. Representative trajectory of *hb-MS2* was shown.

**Figure S2.**
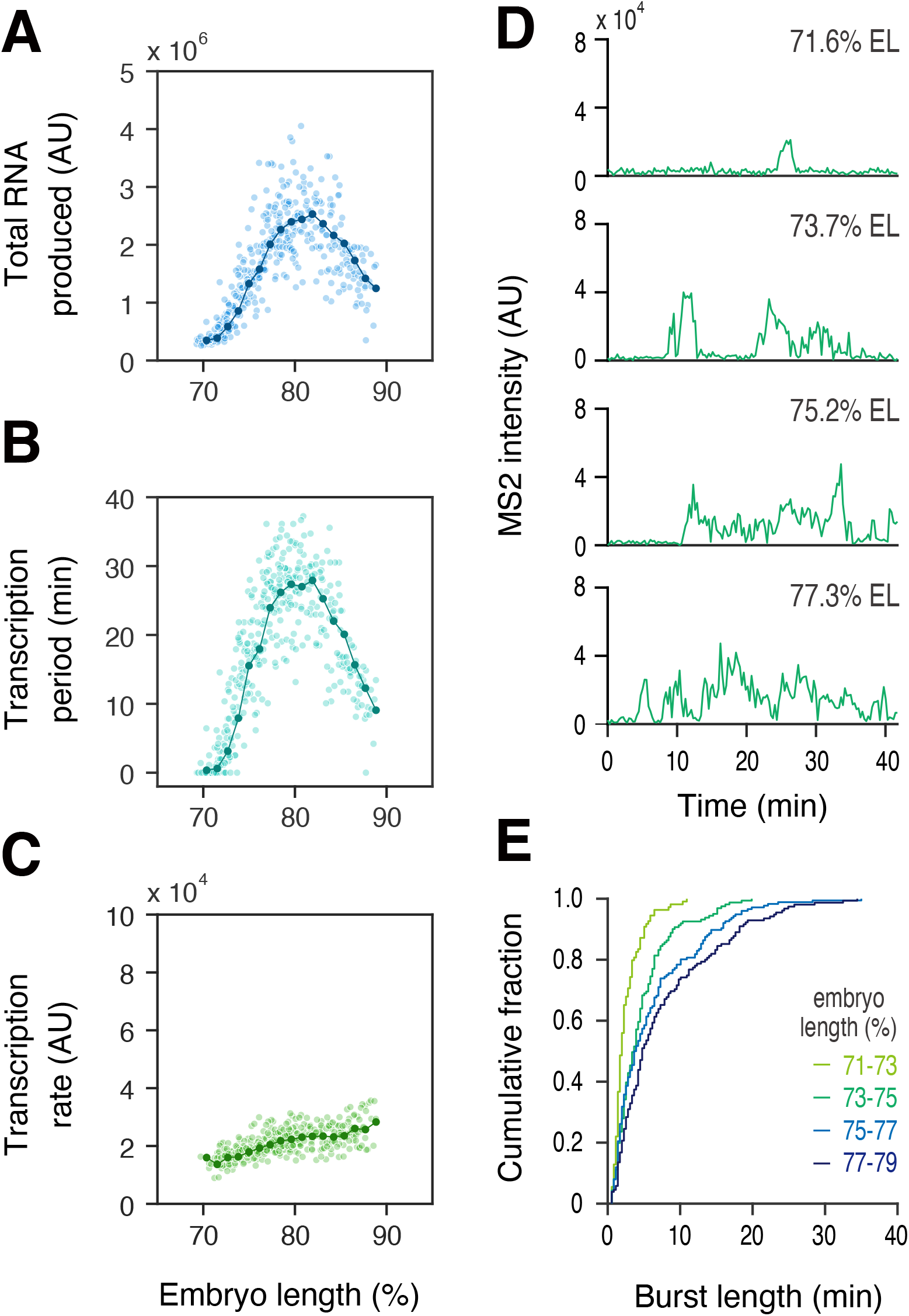
Regulation of *hb* expression at the posterior part of embryos. (A-C) Profiles of total RNA production (A), transcription period (B) and transcription rate (C) as a function of AP position. Line plot indicates mean values in groups of nuclei binned by AP position. In total, 429 nuclei from three independent embryos were analyzed. (D) Representative trajectories of transcription activity of *hb-MS2* in individual nuclei. (E) A cumulative plot showing fraction of bursting events (y axis) and length of transcriptional bursting (x axis). A total of 217 nuclei from three independent embryos was analyzed.

**Figure S3.**
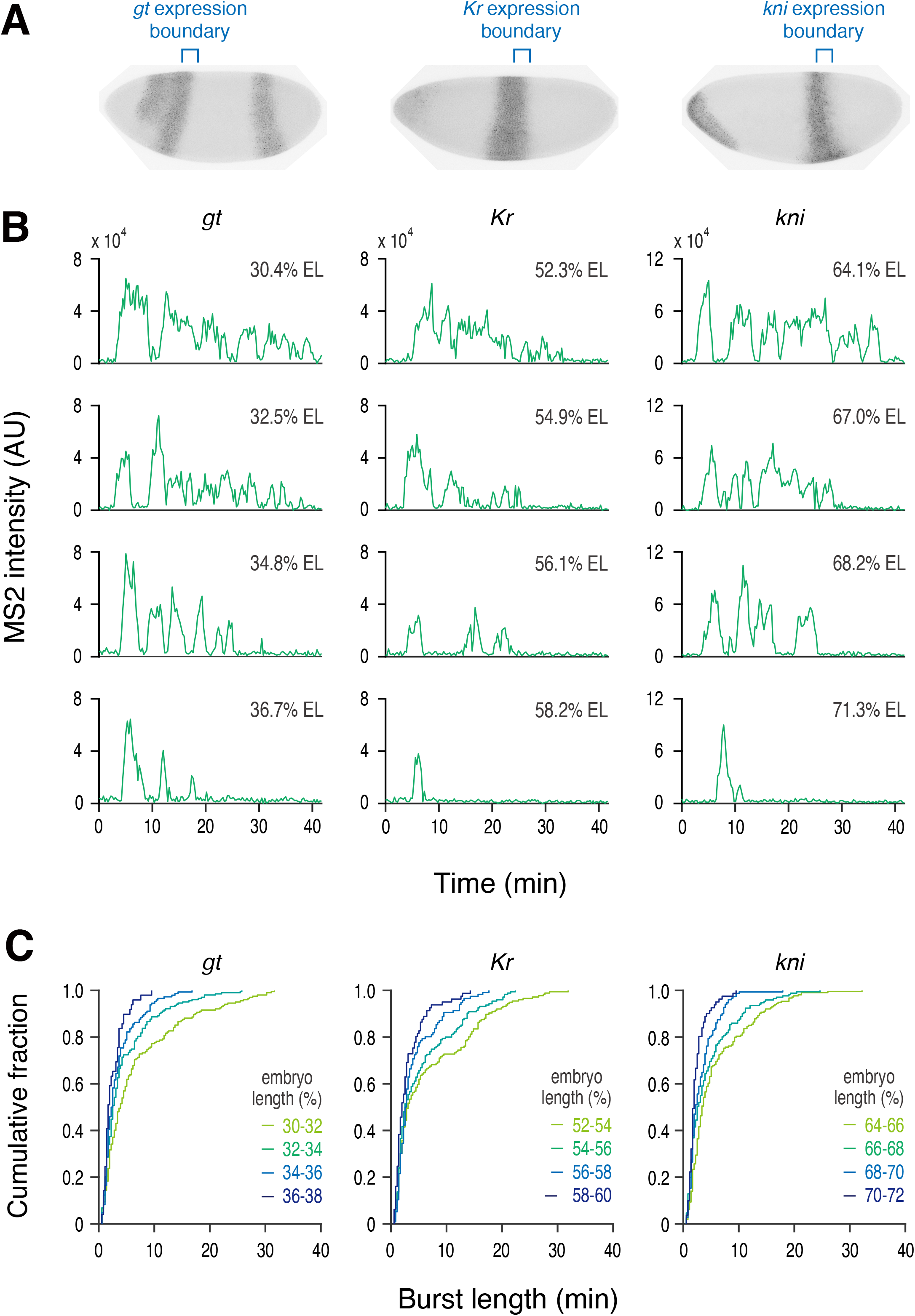
Regulation of transcriptional bursting at expression boundaries. (A) Nuclei surrounding posterior expression boundary were analyzed. Images of *gt, Kr and kni* are same as the images in Figure S1A. (B) Representative trajectories of transcription activity of *gt-MS2* (left), *Kr-MS2* (middle) and *kni-MS2* (right) in individual nuclei. (C) Cumulative plot showing fraction of bursting events (y axis) and length of transcriptional bursting (x axis) from the analysis of three independent embryos. A total of 196, 233, 216 nuclei, respectively, was analyzed from three independent embryos for *gt*, *Kr* and *kni*.

**Figure S4.**
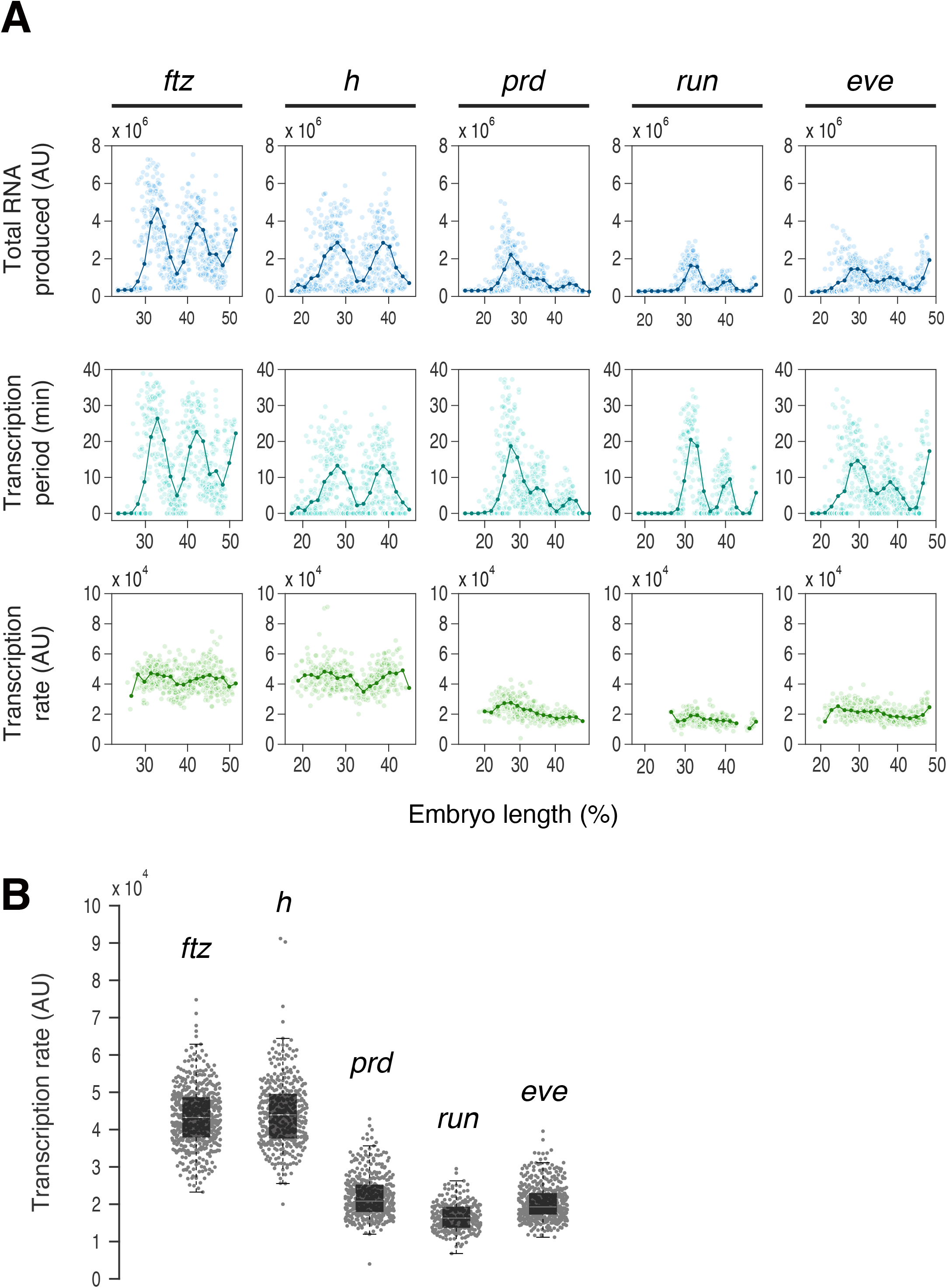
Regulation of pair-rules genes at the stripe1 and stripe2 region. (A) Profiles of total RNA production (top), transcription period (middle) and transcription rate (bottom) as a function of AP position. Line plot indicates mean values in groups of nuclei binned by AP position. A total of 656, 571, 638, 630, 671 nuclei, respectively, were analyzed from three independent embryos for *ftz*, *h*, *prd*, *run*, and *eve*. (B) Boxplot showing the distribution of transcription rate shown in (A). The box indicates the lower (25%) and upper (75%) quantiles, and the solid line indicates the median. Whiskers extend to the most extreme, non-outlier data points.

**Movie S1. Live imaging of***hb-MS2* **at the anterior expression domain.**

Live imaging of *hb-MS2* at the anterior part of embryo during nc14. Each nucleus was false-colored with the intensity proportional to the instantaneous MS2 signal at given time in a given nucleus. The maximum projected image of His2Av-mRFP is shown in gray. Image is oriented with anterior to the left.

**Movie S2. Live imaging of***hb-MS2* **at the posterior expression domain.**

Live imaging of *hb-MS2* at the posterior part of embryo during nc14. Each nucleus was false-colored with the intensity proportional to the instantaneous MS2 signal at given time in a given nucleus. The maximum projected image of His2Av-mRFP is shown in gray. Image is oriented with anterior to the left.

**Movie S3. Live imaging of WT and mutant***hb-MS2* **at the anterior expression domain.**

Live imaging of WT (top) and Δshadow enhancer mutant *hb-MS2* (bottom) at the anterior part of embryo during nc14. Each nucleus was false-colored with the intensity proportional to the instantaneous MS2 signal at given time in a given nucleus. The maximum projected image of His2Av-mRFP is shown in gray. Images are oriented with anterior to the left.

## References

1. Nusslein-Volhard, C., and Wieschaus, E. (1980). Mutations affecting segment number and polarity in *Drosophila*. Nature 287, 795–801.

2. Wakimoto, B.T., and Kaufman, T.C. (1981). Analysis of larval segmentation in lethal genotypes associated with the antennapedia gene complex in *Drosophila melanogaster*. Dev Biol 81, 51–64.

3. Jurgens, G., Wieschaus, E., Nusslein-Volhard, C., and Kluding, H. (1984). Mutations affecting the pattern of the larval cuticle in *Drosophila* melanogaster : II. Zygotic loci on the third chromosome. Wilehm Roux Arch Dev Biol 193, 283–295.

4. Nusslein-Volhard, C., Wieschaus, E., and Kluding, H. (1984). Mutations affecting the pattern of the larval cuticle in *Drosophila* melanogaster : I. Zygotic loci on the second chromosome. Wilehm Roux Arch Dev Biol 193, 267–282.

5. Wieschaus, E., Nusslein-Volhard, C., and Jurgens, G. (1984). Mutations affecting the pattern of the larval cuticle in *Drosophila* melanogaster : III. Zygotic loci on the X-chromosome and fourth chromosome. Wilehm Roux Arch Dev Biol 193, 296–307.

6. Jaeger, J. (2011). The gap gene network. Cell Mol Life Sci 68, 243–274.

7. Jaeger, J., Surkova, S., Blagov, M., Janssens, H., Kosman, D., Kozlov, K.N., Manu, Myasnikova, E., Vanario-Alonso, C.E., Samsonova, M., et al. (2004). Dynamic control of positional information in the early *Drosophila* embryo. Nature 430, 368–371.

8. Bothma, J.P., Garcia, H.G., Esposito, E., Schlissel, G., Gregor, T., and Levine, M. (2014). Dynamic regulation of *eve* stripe 2 expression reveals transcriptional bursts in living *Drosophila* embryos. Proc Natl Acad Sci U S A 111, 10598–10603.

9. Chong, S., Chen, C., Ge, H., and Xie, X.S. (2014). Mechanism of transcriptional bursting in bacteria. Cell 158, 314–326.

10. Chubb, J.R., Trcek, T., Shenoy, S.M., and Singer, R.H. (2006). Transcriptional pulsing of a developmental gene. Curr Biol 16, 1018–1025.

11. Larson, D.R., Zenklusen, D., Wu, B., Chao, J.A., and Singer, R.H. (2011). Real-time observation of transcription initiation and elongation on an endogenous yeast gene. Science 332, 475–478.

12. Little, S.C., Tikhonov, M., and Gregor, T. (2013). Precise developmental gene expression arises from globally stochastic transcriptional activity. Cell 154, 789–800.

13. Pare, A., Lemons, D., Kosman, D., Beaver, W., Freund, Y., and McGinnis, W. (2009). Visualization of individual *Scr* mRNAs during *Drosophila* embryogenesis yields evidence for transcriptional bursting. Curr Biol 19, 2037–2042.

14. Raj, A., Peskin, C.S., Tranchina, D., Vargas, D.Y., and Tyagi, S. (2006). Stochastic mRNA synthesis in mammalian cells. PLoS Biol 4, e309.

15. Suter, D.M., Molina, N., Gatfield, D., Schneider, K., Schibler, U., and Naef, F. (2011). Mammalian genes are transcribed with widely different bursting kinetics. Science 332, 472–474.

16. Zenklusen, D., Larson, D.R., and Singer, R.H. (2008). Single-RNA counting reveals alternative modes of gene expression in yeast. Nat Struct Mol Biol 15, 1263–1271.

17. Dubuis, J.O., Samanta, R., and Gregor, T. (2013). Accurate measurements of dynamics and reproducibility in small genetic networks. Mol Syst Biol 9, 639.

18. Garcia, H.G., Tikhonov, M., Lin, A., and Gregor, T. (2013). Quantitative imaging of transcription in living *Drosophila* embryos links polymerase activity to patterning. Curr Biol 23, 2140–2145.

19. Lucas, T., Ferraro, T., Roelens, B., De Las Heras Chanes, J., Walczak, A.M., Coppey, M., and Dostatni, N. (2013). Live imaging of bicoid-dependent transcription in *Drosophila* embryos. Curr Biol 23, 2135–2139.

20. Jinek, M., East, A., Cheng, A., Lin, S., Ma, E., and Doudna, J. (2013). RNA-programmed genome editing in human cells. Elife 2, e00471.

21. Harding, K., Hoey, T., Warrior, R., and Levine, M. (1989). Autoregulatory and gap gene response elements of the *even-skipped* promoter of *Drosophila*. EMBO J 8, 1205–1212.

22. Jiang, J., Hoey, T., and Levine, M. (1991). Autoregulation of a segmentation gene in *Drosophila*: combinatorial interaction of the *even-skipped* homeo box protein with a distal enhancer element. Genes Dev 5, 265–277.

23. Schier, A.F., and Gehring, W.J. (1992). Direct homeodomain-DNA interaction in the autoregulation of the fushi tarazu gene. Nature 356, 804–807.

24. Struhl, G., Struhl, K., and Macdonald, P.M. (1989). The gradient morphogen bicoid is a concentration-dependent transcriptional activator. Cell 57, 1259–1273.

25. Wimmer, E.A., Carleton, A., Harjes, P., Turner, T., and Desplan, C. (2000). Bicoid-independent formation of thoracic segments in *Drosophila*. Science 287, 2476–2479.

26. Margolis, J.S., Borowsky, M.L., Steingrimsson, E., Shim, C.W., Lengyel, J.A., and Posakony, J.W. (1995). Posterior stripe expression of hunchback is driven from two promoters by a common enhancer element. Development 121, 3067–3077.

27. Perry, M.W., Boettiger, A.N., and Levine, M. (2011). Multiple enhancers ensure precision of gap gene-expression patterns in the *Drosophila* embryo. Proc Natl Acad Sci U S A 108, 13570–13575.

28. Frankel, N., Davis, G.K., Vargas, D., Wang, S., Payre, F., and Stern, D.L. (2010). Phenotypic robustness conferred by apparently redundant transcriptional enhancers. Nature 466, 490–493.

29. Perry, M.W., Boettiger, A.N., Bothma, J.P., and Levine, M. (2010). Shadow enhancers foster robustness of *Drosophila* gastrulation. Curr Biol 20, 1562–1567.

30. Bothma, J.P., Garcia, H.G., Ng, S., Perry, M.W., Gregor, T., and Levine, M. (2015). Enhancer additivity and non-additivity are determined by enhancer strength in the *Drosophila* embryo. Elife 4.

31. Driever, W., and Nusslein-Volhard, C. (1989). The bicoid protein is a positive regulator of *hunchback* transcription in the early *Drosophila* embryo. Nature 337, 138–143.

32. Wu, H., Manu, Jiao, R., and Ma, J. (2015). Temporal and spatial dynamics of scaling-specific features of a gene regulatory network in *Drosophila*. Nat Commun 6, 10031.

33. Boettiger, A.N., and Levine, M. (2013). Rapid transcription fosters coordinate *snail* expression in the *Drosophila* embryo. Cell Rep 3, 8–15.

34. Edgar, B.A., Weir, M.P., Schubiger, G., and Kornberg, T. (1986). Repression and turnover pattern *fushi tarazu* RNA in the early *Drosophila* embryo. Cell 47, 747–754.

35. Bartman, C.R., Hsu, S.C., Hsiung, C.C., Raj, A., and Blobel, G.A. (2016). Enhancer Regulation of Transcriptional Bursting Parameters Revealed by Forced Chromatin Looping. Mol Cell 62, 237–247.

36. Fukaya, T., Lim, B., and Levine, M. (2016). Enhancer Control of Transcriptional Bursting. Cell 166, 358–368.

37. Boija, A., Klein, I.A., Sabari, B.R., Dall’Agnese, A., Coffey, E.L., Zamudio, A.V., Li, C.H., Shrinivas, K., Manteiga, J.C., Hannett, N.M., et al. (2018). Transcription Factors Activate Genes through the Phase-Separation Capacity of Their Activation Domains. Cell 175, 1842–1855 e1816.

38. Cho, W.K., Spille, J.H., Hecht, M., Lee, C., Li, C., Grube, V., and Cisse, II (2018). Mediator and RNA polymerase II clusters associate in transcription-dependent condensates. Science 361, 412–415.

39. Sabari, B.R., Dall’Agnese, A., Boija, A., Klein, I.A., Coffey, E.L., Shrinivas, K., Abraham, B.J., Hannett, N.M., Zamudio, A.V., Manteiga, J.C., et al. (2018). Coactivator condensation at super-enhancers links phase separation and gene control. Science 361.

40. Dunipace, L., Ozdemir, A., and Stathopoulos, A. (2011). Complex interactions between cis-regulatory modules in native conformation are critical for *Drosophila snail* expression. Development 138, 4075–4084.

41. Perry, M.W., Bothma, J.P., Luu, R.D., and Levine, M. (2012). Precision of hunchback expression in the *Drosophila* embryo. Curr Biol 22, 2247–2252.

42. Antolovic, V., Miermont, A., Corrigan, A.M., and Chubb, J.R. (2017). Generation of Single-Cell Transcript Variability by Repression. Curr Biol 27, 1811–1817 e1813.

43. Zoller, B., Little, S.C., and Gregor, T. (2018). Diverse Spatial Expression Patterns Emerge from Unified Kinetics of Transcriptional Bursting. Cell 175, 835–847 e825.

44. Yokoshi, M., Segawa, K., and Fukaya, T. (2020). Visualizing the Role of Boundary Elements in Enhancer-Promoter Communication. Mol Cell 78, 224–235 e225.

45. Lim, B., Fukaya, T., Heist, T., and Levine, M. (2018). Temporal dynamics of pair-rule stripes in living *Drosophila* embryos. Proc Natl Acad Sci U S A 115, 8376–8381.

46. Ren, X.J., Sun, J., Housden, B.E., Hu, Y.H., Roesel, C., Lin, S.L., Liu, L.P., Yang, Z.H., Mao, D.C., Sun, L.Z., et al. (2013). Optimized gene editing technology for *Drosophila melanogaster* using germ line-specific Cas9. P Natl Acad Sci USA 110, 19012–19017.

47. Ringrose, L. (2009). Transgenesis in *Drosophila* melanogaster. Methods Mol Biol 561, 3–19.

48. Dufourt, J., Trullo, A., Hunter, J., Fernandez, C., Lazaro, J., Dejean, M., Morales, L., Nait-Amer, S., Schulz, K.N., Harrison, M.M., et al. (2018). Temporal control of gene expression by the pioneer factor Zelda through transient interactions in hubs. Nat Commun 9.

49. Lim, B., Heist, T., Levine, M., and Fukaya, T. (2018). Visualization of Transvection in Living *Drosophila* Embryos. Mol Cell 70, 287–296 e286.

50. Li, X.Y., Harrison, M.M., Villalta, J.E., Kaplan, T., and Eisen, M.B. (2014). Establishment of regions of genomic activity during the *Drosophila* maternal to zygotic transition. Elife 3.

51. Koenecke, N., Johnston, J., Gaertner, B., Natarajan, M., and Zeitlinger, J. (2016). Genome-wide identification of *Drosophila* dorso-ventral enhancers by differential histone acetylation analysis. Genome Biol 17, 196.

52. Wang, Y.L., Duttke, S.H., Chen, K., Johnston, J., Kassavetis, G.A., Zeitlinger, J., and Kadonaga, J.T. (2014). TRF2, but not TBP, mediates the transcription of ribosomal protein genes. Genes Dev 28, 1550–1555.

